# Mantis-Delta: Mass-Action Network Theory and Steady-State Characterization for Chemical Reaction Networks

**DOI:** 10.64898/2026.05.14.725189

**Authors:** Emilio A. Venegas

## Abstract

Chemical Reaction Network Theory (CRNT), developed by Horn, Jackson, and Feinberg, provides parameter-free structural theorems that constrain the asymptotic dynamics of mass-action systems irrespective of the numerical values of the rate constants. Despite the maturity of the theory, modern open-source implementations that combine CRNT structural analysis with symbolic ordinary differential equation (ODE) construction and robust numerical steady-state finding remain scarce. We present `mantis-delta`, a pure Python library that ingests human-readable reaction strings, builds the complex reaction graph, computes the deficiency *δ* = *n*−*ℓ*−*s* and weak reversibility, and decides applicability of the Deficiency Zero Theorem (DZT) and Deficiency One Theorem (D1T). For systems satisfying these structural conditions, `mantis-delta` certifies, without any simulation whatsoever, existence, uniqueness and (for DZT) asymptotic stability of the positive steady state in every stoichiometric compatibility class. When the structural theorems do not apply, the library provides symbolic mass-action ODEs and Jacobians via SymPy and a hybrid numerical solver that combines stiff implicit integration with bound-constrained algebraic least-squares to locate both stable and unstable fixed points, including Hopf bifurcation centres inaccessible to forward integration. We demonstrate the workflow on six benchmarks: a reversible isomerisation, the Michaelis–Menten enzyme mechanism, the closed and chemostatted Brusselator, a catalytic hairpin assembly (CHA) miR-21 biosensor, and the Goldbeter–Koshland zero-order ultrasensitivity switch. In each case, the CRNT-predicted qualitative behaviour (monostability, oscillation, uniqueness) is recovered numerically with a residual below 10^−6^ M s^−1^, and the Goldbeter–Koshland dose-response curve agrees with the closed-form quasi-steady-state approximation to within 1% over a 400× kinase/phosphatase activity scan. `mantis-delta` is open-source (MIT license) and available at https://github.com/emiliovenegas/mantis-delta.

## 1 Introduction

The dynamics of a chemical reaction network (CRN) endowed with mass-action kinetics are governed by polynomial ordinary differential equations whose behaviour, *a priori*, depends jointly on the network topology and on *O*(10^0^–10^2^) kinetic parameters that are seldom known to better than one order of magnitude in living systems. A central insight of twentieth-century mathematical chemistry, originating in the work of Horn, Jackson, and Feinberg, is that a substantial portion of the qualitative behaviour, existence and uniqueness of equilibria, monostability, impossibility of oscillation, is dictated by *purely structural* invariants of the reaction graph (Horn and Jackson 1972; Feinberg 1972, 1987). The two cornerstones of this body of results are the Deficiency Zero Theorem (Horn and Jackson 1972; Feinberg 1987) and the Deficiency One Theorem (Feinberg 1995), each of which guarantees properties of the steady-state set that hold uniformly over *every* assignment of positive rate constants. Modern monographs (Feinberg 2019) and review articles (Angeli 2009) provide a complete account of the theory, and recent results on the global attractor conjecture (Anderson 2011) and structural sources of robustness (Shinar and Feinberg 2010) continue to extend its reach.

Notwithstanding the maturity of CRNT, the practical bridge between formal theorems and computational pipelines remains under-served. The Chemical Reaction Network Toolbox (Ji et al. 2011) and the constraint-programming approach of Soliman (2012) implement parts of the deficiency machinery, but neither provides an end-to-end Python interface that (i) accepts the reaction strings biochemists actually write, (ii) exposes the mass-action ODEs and Jacobian symbolically through SymPy (Meurer et al. 2017), and (iii) couples the structural analysis to a robust numerical solver capable of finding both stable and *unstable* fixed points, the latter being indispensable for the location of Hopf bifurcation centres in chemostatted, open systems (Prigogine and Lefever 1968).

In this paper we introduce `mantis-delta`, a small Python library that fills this gap. Built on top of the SciPy scientific stack (Virtanen et al. 2020; Harris et al. 2020), SymPy (Meurer et al. 2017), and NetworkX (Hagberg et al. 2008), `mantis-delta` implements:

1. parsing of human-readable reaction strings, canonical-form normalisation of rate-constant keys, and construction of the complex reaction graph;
2. exact rational computation of the stoichiometric matrix rank, deficiency *δ*, linkage classes, weak reversibility, and conservation laws as non-negative integer left null-space vectors;
3. decision procedures for the Deficiency Zero and Deficiency One Theorems, including the per-linkage-class deficiency checks required by D1T (Feinberg 1995, 2019);
4. symbolic mass-action ODE and Jacobian construction with both symbolic and numeric rate constants;
5. a hybrid steady-state solver combining the stiff implicit Radau integrator (Hairer and Wanner 1996; Hindmarsh et al. 2005) with bound-constrained algebraic least-squares;
6. a wrapper for chemostatted (open) networks in the sense of Polettini and Esposito (2014), where selected species are held at fixed concentrations and folded into effective rate prefactors;
7. log-spaced bifurcation scans of any rate constant and a NetworkX-based reaction-graph visualiser.

`mantis-delta` targets two distinct user communities: (a) systems biologists and synthetic-biology engineers who want to certify, before any wet-lab work, that an enzymatic cycle (Ferrell and Ha 2014), a DNA strand-displacement cascade (Yin et al. 2008; Li et al. 2011), or a metabolic motif (Ang et al. 2013) cannot exhibit pathological bistability or oscillation; and (b) educators and researchers in chemical reaction network theory who want a compact, fully open-source reference implementation against which to develop further structural algorithms.

The remainder of this article is organised as follows. Section 2 presents the mathematical objects and algorithms implemented in `mantis-delta`. Section 3 applies the library to six benchmark networks of increasing complexity: a reversible isomerisation (DZT applies), Michaelis–Menten (δ = 0 but not weakly reversible), the closed and chemostatted Brusselator (Hopf-driven limit cycle), the catalytic hairpin assembly (CHA) miR-21 biosensor (δ = 1, D1T applies), and the Goldbeter–Koshland zero-order ultrasensitivity switch (Goldbeter and Koshland 1981). Section 4 places `mantis-delta` in the context of existing CRN software and outlines limitations and future directions.

## 2 Mathematical preliminaries and software architecture

### 2.1 Reactions, complexes, and the stoichiometric matrix

We follow the standard notation of Feinberg (2019). Let 𝒮 = {*S*_1_, …, *S*_*m*_} denote the set of *m* chemical species and let 𝒞 denote a finite set of *n complexes*, where a complex 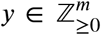 is a non-negative integer linear combination of species. A *reaction* is an ordered pair of distinct complexes, written *y* → *y*′. A reaction network 𝒩 = (𝒮, 𝒞, ℛ) consists of the finite set ℛ of *r* reactions on 𝒞.

The *stoichiometric matrix* **N** ∈ ℤ^*m*×*r*^ has columns indexed by reactions and rows indexed by species, with

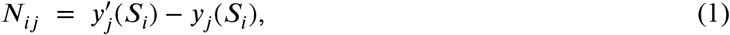

i.e. the net change in copy number of species *i* effected by reaction *j*. The *stoichiometric subspace* is 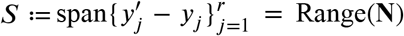, of dimension *s* ≔ rank(**N**). For any trajectory of the kinetic system, the concentration vector 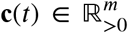 *class* of the initial condition **c**_0_. is confined to the affine variety **c**_0_ + *S*, the *stoichiometric compatibility*

### 2.2 Mass-action kinetics

Under the law of mass action (Horn and Jackson 1972; Michaelis and Menten 1913), the flux of the *j*-th reaction 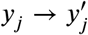 with rate constant *k*_*j*_ > 0 is

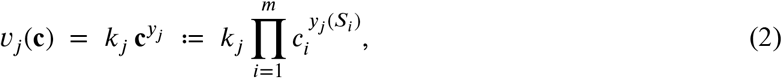

and the species-concentration ODE system reads

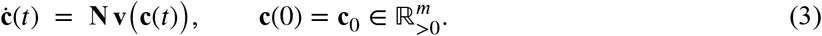

The Jacobian along a trajectory is

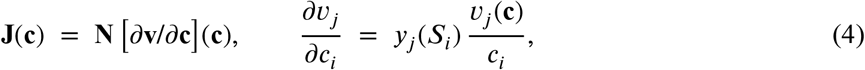

with the convention that the right-hand side equals 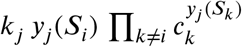 when *c*_*i*_ = 0. In `mantis-delta`, equations (Equation 3)–(Equation 4) are constructed symbolically as SymPy expressions via `network.CRNetwork.odes()` and `network.CRNetwork.jacobian()`; rate constants may be kept symbolic (*k*_1_, *k*_2_, …) or substituted by their numeric values.

### 2.3 Conservation laws

Conservation laws are vectors 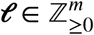 in the *left* null space of **N**,

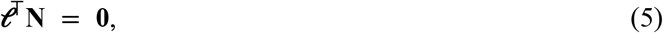

so that 𝓁^⊤^**c**(*t*) is invariant along every trajectory. `mantis-delta` computes a non-negative integer basis of LeftNull(**N**) via exact rational arithmetic with SymPy, then post-processes the basis (sign normalisation, integer scaling, and elementary reduction) to recover physical moiety totals. The algorithm exploits the fact that, for mass-balanced networks, a non-negative integer basis is always admissible (Feinberg 2019; Rao et al. 2013).

### 2.4 Deficiency, linkage classes, and weak reversibility

The *reaction graph G* = (𝒞, ℛ) has the complexes as nodes and the reactions as directed edges. A *linkage class* is a weakly connected component of *G*; we denote their number by 𝓁. The *deficiency* of the network is

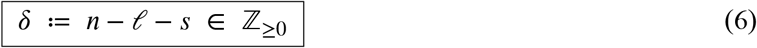

where *n* = |𝒞 |, 𝓁 is the number of linkage classes, and *s* = rank(**N**). The non-negativity of *δ* is a classical fact (Feinberg 1987). The network is *weakly reversible* if every weakly connected component of *G* is also strongly connected.

### 2.5 Feinberg’s structural theorems

The two principal results implemented in `mantis-delta` are:

#### Theorem 1

**(Deficiency Zero Theorem; Horn–Jackson 1972, Feinberg 1972)**. *Let* 𝒩 *be a weakly reversible reaction network with δ* = 0. *Then for every choice of positive rate constants* 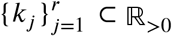, *the mass-action system (Equation 3) admits exactly one positive equilibrium in every positive stoichiometric compatibility class, and that equilibrium is locally asymptotically stable. In particular, sustained oscillations and multistability are impossible* (Horn and Jackson 1972; Feinberg 2019).

#### Theorem 2

**(Deficiency One Theorem; Feinberg 1995)**. *Let* 𝒩 *have deficiency δ* = 1 *and assume that each linkage class L*_*q*_ *has per-linkage-class deficiency δ*_*q*_ ∈ {0, 1} *with at most one δ*_*q*_ = 1. *Then for every choice of positive rate constants, the mass-action system (Equation 3) admits at most one positive equilibrium in every positive stoichiometric compatibility class* (Feinberg 1995, 2019).

`mantis-delta` decides the structural hypotheses of both theorems by direct graph construction (NetworkX), exact rank computation (NumPy (Harris et al. 2020)), and per-LC deficiency calculation via QR-pivoted row selection of **N** (Virtanen et al. 2020). The decision is returned in human-readable form by `CRNetwork.crnt_summary()` and programmatically by the `CRNTResult` dataclass.

### 2.6 Open (chemostatted) networks

A *chemostatted* species is one whose concentration is held constant by an external reservoir, mimicking continuous replenishment in a flow reactor or a buffer with effectively infinite capacity. Following the open-network formalism of Polettini and Esposito (2014), `mantis-delta` removes chemostatted species from the dynamic state vector and folds their (fixed) concentrations into *effective* rate constants:

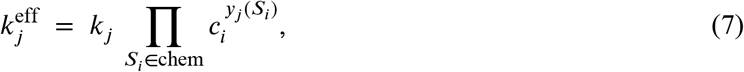

so that the reduced ODE system contains only the dynamic species. Crucially, on open systems the conservation-law left null space is in general empty and forward integration cannot locate Hopf-unstable fixed points; `mantis-delta` therefore switches to a *scale-aware multi-start* bound-constrained least-squares strategy that retains the ability to find unstable equilibria.

### 2.7 Hybrid steady-state solver

For a closed or semi-closed network with *p* independent conservation laws encoded as the rows of **L** ∈ ℤ^*p*×*m*^ and totals 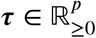, the steady-state problem becomes the constrained polynomial system

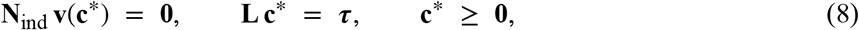

where **N**_ind_ collects *s* linearly independent rows of **N** selected by QR pivoting. `mantis-delta` first attempts forward integration of (Equation 3) with SciPy’s stiff implicit Radau IIA solver (Hairer and Wanner 1996; Virtanen et al. 2020) from the user-supplied initial condition, then runs bound-constrained algebraic least-squares (`scipy.optimize.least_squares`, trust-region reflective method) seeded both at the user IC and at *n*_attempts_ −2 random points on the conservation manifold. For open networks the integration step is skipped and only the algebraic strategy is used. Duplicate solutions (relative 𝓁_2_ distance below 1%) and non-physical solutions are filtered, after which the Jacobian (Equation 4) is evaluated at each candidate and its eigenvalues classified as in Table 1.

**Table 1:**
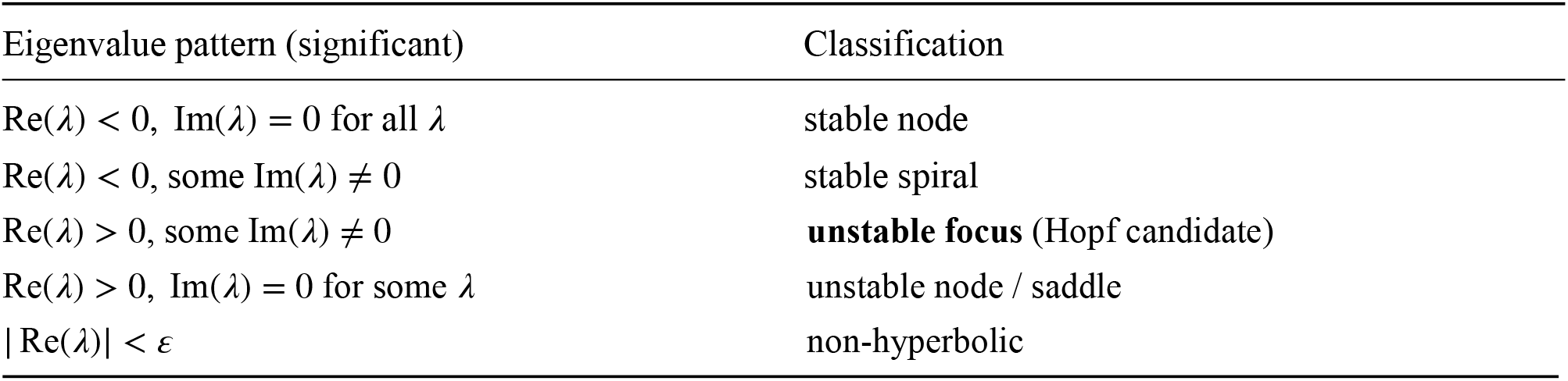
Stability classification implemented in analysis.classify_steady_state(). Eigenvalues with |*λ*| < 10^−4^ max_*k*_ |*λ*_*k*_| are treated as zero (artefacts of the conservation-law dimensions) and excluded. The `is_oscillatory` flag is set whenever any significant eigenvalue has a non-trivial imaginary part, covering both stable spirals and unstable foci.

## 3 Results

We illustrate `mantis-delta`’s end-to-end workflow on six benchmark networks of increasing complexity. All examples are bundled with the source distribution (`examples/01–06`).

### 3.1 Reversible isomerisation: parameter-free uniqueness

The simplest non-trivial network is the reversible isomerisation *A* ⇌ *B*. There are *n* = 2 complexes, 𝓁 = 1 linkage class, rank(**N**) = 1, hence *δ* = 0. The network is weakly reversible, so the Deficiency Zero Theorem applies; for any positive forward and reverse rate constants *k*_*f*_, *k*_*r*_ the unique positive equilibrium consistent with the conservation law *A* + *B* = *T* is

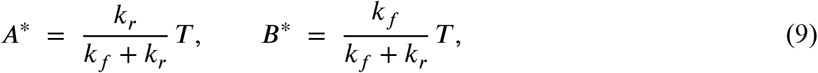

and it is asymptotically stable. `mantis-delta` recovers (Equation 9) numerically to machine precision: for *k*_*f*_ = 1.0, *k*_*r*_ = 0.5, *A*_0_ = 2, *B*_0_ = 0 the solver returns 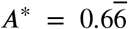 and 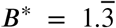 with residual ‖**Nv**(**c**^∗^)‖_2_ < 10^−12^ M s^−1^ and eigenvalues {0, −1.5}. The single zero eigenvalue is the expected signature of the conservation law dimension.

The network is constructed from a human-readable string via `CRNetwork.from_string`; `crnt_summary` reports *δ* = 0 and confirms weak reversibility, enabling the DZT; and `steady_states` locates the fixed point and returns both the equilibrium concentrations and the eigenvalues of the Jacobian.

### 3.2 Michaelis–Menten: *δ* = 0 but not weakly reversible

The classical enzyme mechanism (Michaelis and Menten 1913)

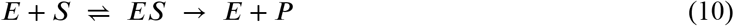

has three complexes, one linkage class, and rank(**N**) = 2, hence *δ* = 0. However the network is *not* weakly reversible because the catalytic step *ES* → *E* +*P* has no reverse edge in the complex graph. The Deficiency Zero Theorem therefore *does not* apply in its stability form. `mantis-delta` correctly reports this and returns the two conservation laws *E* + *ES* = *E*_tot_ and *S* + *ES* + *P* = *S*_tot_ as non-negative integer left null-space vectors. For the standard parameters *k*_*f*_ = 10^6^ M^−1^s^−1^, *k*_*r*_ = 10^3^ s^−1^, *k*_cat_ = 100 s^−1^ and initial conditions *E*_0_ = 1 *μ*M, *S*_0_ = 100 *μ*M, the numerical equilibrium satisfies enzyme conservation *E*^∗^ + *ES*^∗^ = *E*_0_ to within machine precision, and the quasi-steady-state estimate

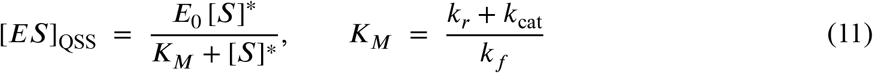

tracks the full mass-action concentration [*ES*](*t*) obtained by `mantis-delta` to better than 0.25% throughout the catalytic phase (Fig. 1). At the true mathematical steady state the irreversible catalytic step drives all substrate to product, so [*ES*]^∗^ = 0; the practical relevance of the QSS approximation is therefore as a *dynamic* tracker during enzyme turnover rather than as a steady-state identity.

**Figure 1:**
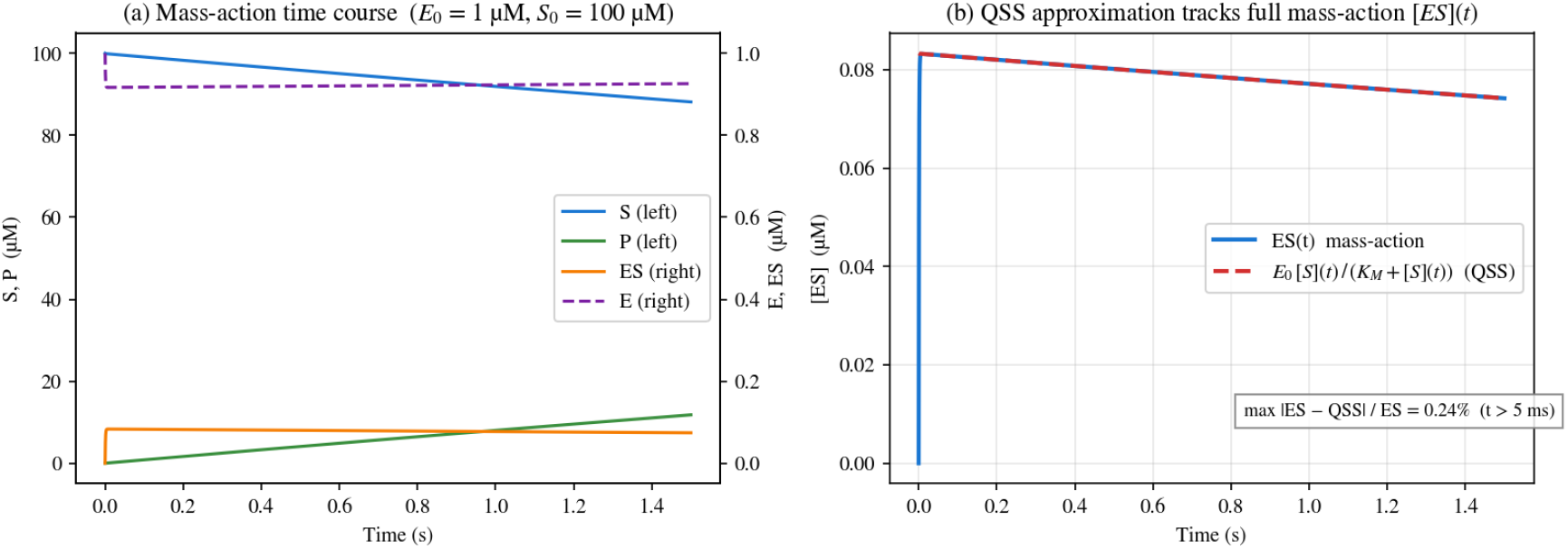
Michaelis–Menten enzyme kinetics. (a) Full mass-action time course (*E*_0_ = 1 *μ*M, *S*_0_ = 100 *μ*M) on a twin-axis plot: *S* and *P* on the left axis, *E* and *ES* on the right. (b) Comparison of the full mass-action [*ES*](*t*) (solid) with the quasi-steady-state approximation *E*_0_ [*S*](*t*)/(*K*_*M*_ + [*S*](*t*)) (dashed); the two curves overlap to within 0.25% after the initial ~ 5 ms transient. Generated by `examples/02_michaelis_menten.py`.

**Figure 2:**
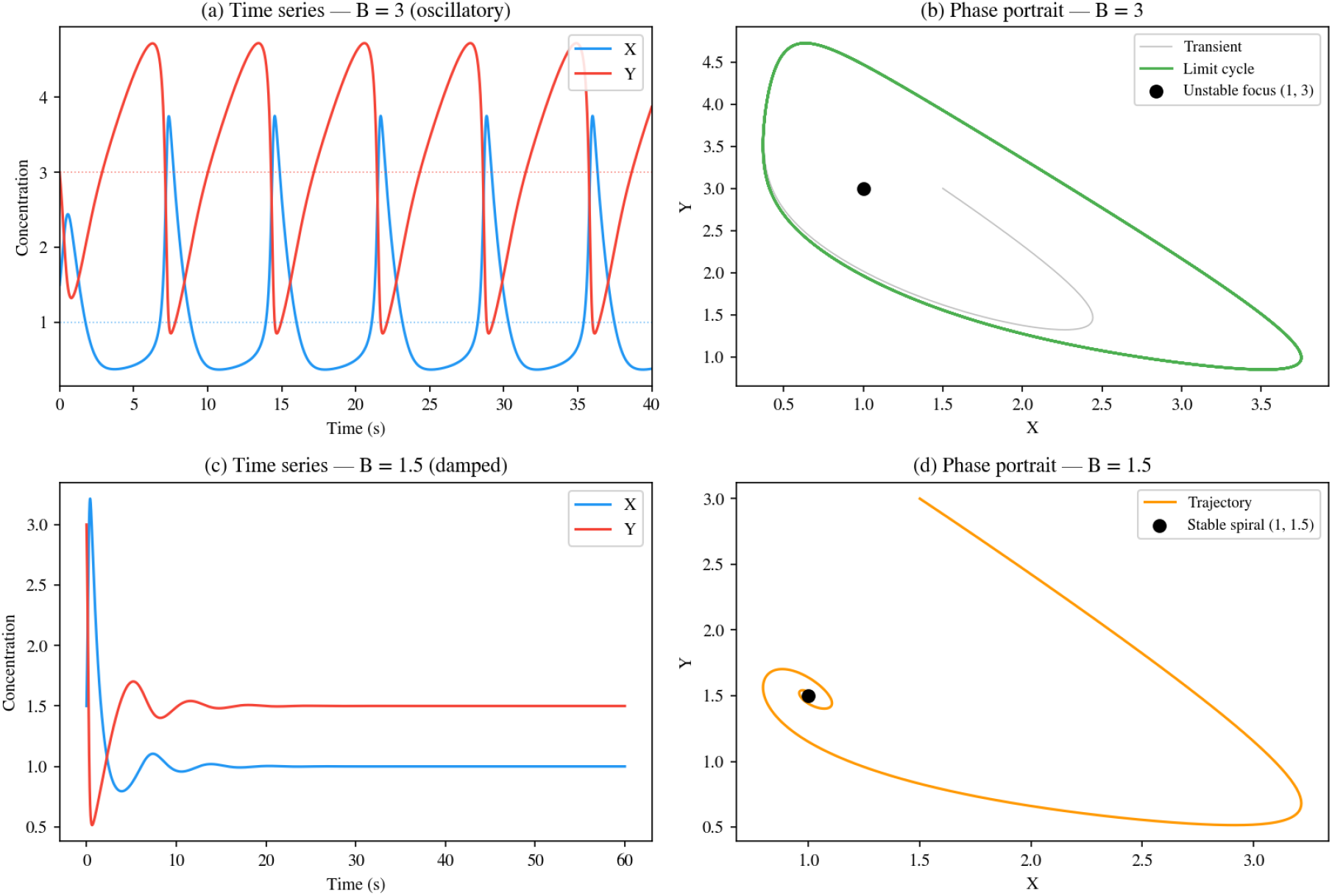
Brusselator dynamics with *A* and *B* chemostatted. Top row (*B* = 3, oscillatory): (a) time series of *X* and *Y*; (b) phase portrait showing the transient (grey) relaxing onto the limit cycle (green) that orbits the algebraic fixed point (*X*^∗^, *Y* ^∗^) = (1, 3), classified as an unstable focus by `mantis-delta`. Bottom row (*B* = 1.5, damped): (c) time series; (d) phase portrait spiralling into the stable equilibrium (1, 1.5). The black markers indicate the equilibria returned by `mantis-delta`’s algebraic solver. Generated by `examples/05_brusselator_chemostatted.py`.

### 3.3 Brusselator: Hopf bifurcation under chemostatting

The four-step Brusselator (Prigogine and Lefever 1968)

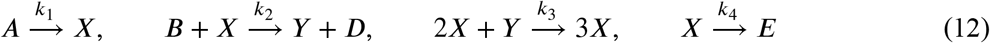

is the canonical example of a chemical oscillator. With *all* six species treated as dynamic, `mantis-delta` reports *δ* = 0 but the network is not weakly reversible (no autocatalytic complex appears as a product of any other reaction), so neither Feinberg theorem certifies global asymptotic stability. Forward integration of the full closed system converges to a fixed point because *A* and *B* deplete; this is the well-known distinction between the *closed* and the *classical chemostatted* Brusselator (Prigogine and Lefever 1968).

Holding *A* and *B* at constant concentrations *A* = 1, *B* = 3 via the `chemostatted=` argument folds their values into effective rate constants (Equation 7) and reduces the system to the standard two-variable form

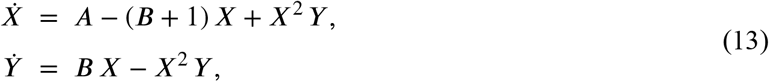

with unique fixed point *X*^∗^ = *A, Y* ^∗^ = *B*/*A*. Linearisation yields the Jacobian

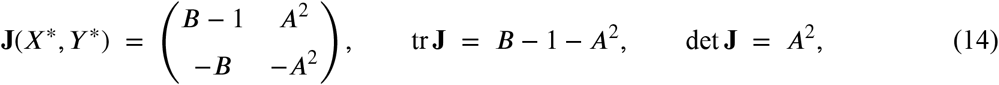

so that a supercritical Hopf bifurcation occurs at the threshold *B* = 1 + *A*^2^ (Prigogine and Lefever 1968). For *A* = 1, *B* = 3 the condition *B* > 1 + *A*^2^ is satisfied and `mantis-delta` locates the *unstable* fixed point (*X*^∗^, *Y* ^∗^) = (1, 3) algebraically (forward integration would orbit the limit cycle indefinitely); the Jacobian eigenvalues are 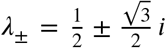 with positive real part, correctly classifying the fixed point as an unstable focus. For *B* = 1.5 < 1 + *A*^2^ the same fixed point becomes a stable spiral.

### 3.4 CHA miR-21 biosensor: parameter-free uniqueness via D1T

DNA strand-displacement networks have emerged as a versatile platform for constructing programmable molecular sensors (Yin et al. 2008; Li et al. 2011). The catalytic hairpin assembly (CHA) circuit for the microRNA biomarker miR-21 comprises four reversible reactions:

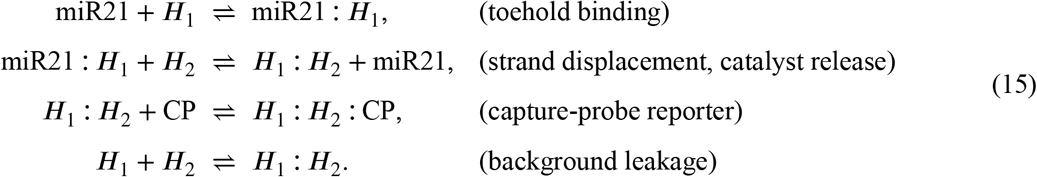

Forward and reverse rate constants were obtained from NUPACK thermodynamic calculations (Zadeh et al. 2011) at 37 °C in 137 mM NaCl, 10 mM MgCl_2_. `mantis-delta`’s structural analysis returns *m* = 7 species, *n* = 8 complexes, 𝓁 = 4 linkage classes, rank(**N**) = 3, hence

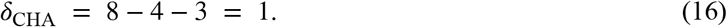

The network is weakly reversible and each linkage class has per-LC deficiency *δ*_*q*_ ∈ {0, 1} with exactly one *δ*_*q*_ = 1, so the Deficiency One Theorem applies and certifies *at most one* positive equilibrium per stoichiometric compatibility class. `mantis-delta` further returns the four moiety totals (miR21_tot_, *H*_1,tot_, *H*_2,tot_, CP_tot_) as a non-negative integer basis of LeftNull(**N**).

With *H*_1,0_ = *H*_2,0_ = CP_0_ = 100 nM and [miR21]_0_ = 10 nM, mass-action integration of the network (Fig. 3) reveals the characteristic catalytic turnover: miR-21 is rapidly consumed into the miR21⋅H_1_ complex, which is then displaced by H_2_ to regenerate free miR-21 and form the H_1_:*H*_2_ dimer that binds the capture probe. The hybrid solver returns the unique equilibrium [*H*_1_:*H*_2_:CP]^∗^ = 0.34 nM, with Re *λ* < 0 for all significant eigenvalues (Re *λ*_max_ ≈ −3 × 10^−3^ s^−1^). A log-spaced scan over [miR21]_0_ ∈ [0.1, 100] nM (Fig. 4) confirms the D1T prediction empirically: the dose-response curve is single-valued and monotone, with no hint of hysteresis or alternative branches.

**Figure 3:**
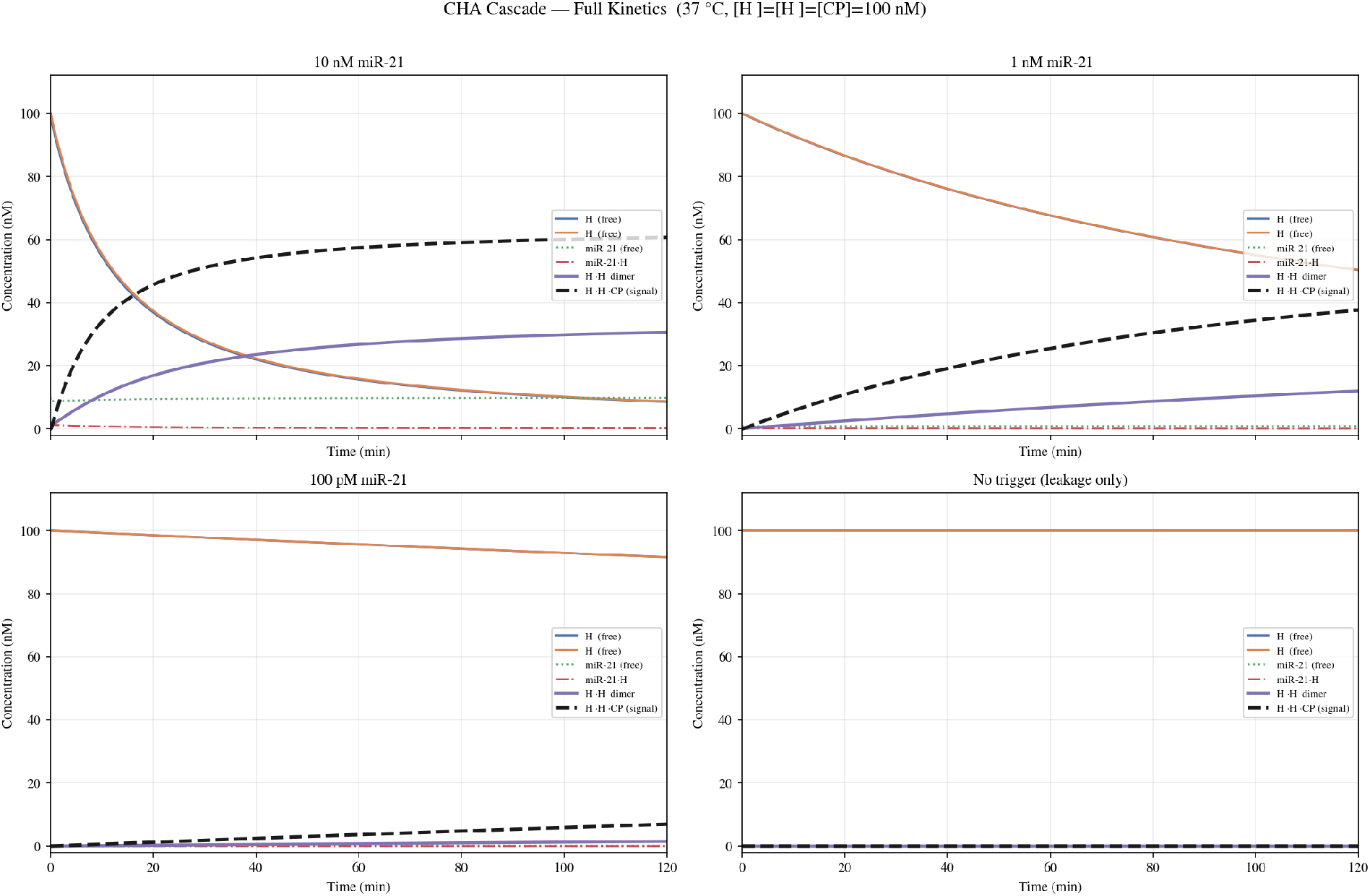
Mass-action kinetics of the CHA cascade at four miR-21 trigger concentrations (10 nM, 1 nM, 100 pM, and no-trigger leakage-only control). The reporter complex *H*_1_:*H*_2_:CP (dashed black) is the electrochemical output; H_1_ and H_2_ are the fuel hairpins held at 100 nM. The leakage-only panel demonstrates that, at the rate constants returned by NUPACK, spontaneous H_1_:*H*_2_ dimerisation is negligible over a 120 min window.

**Figure 4:**
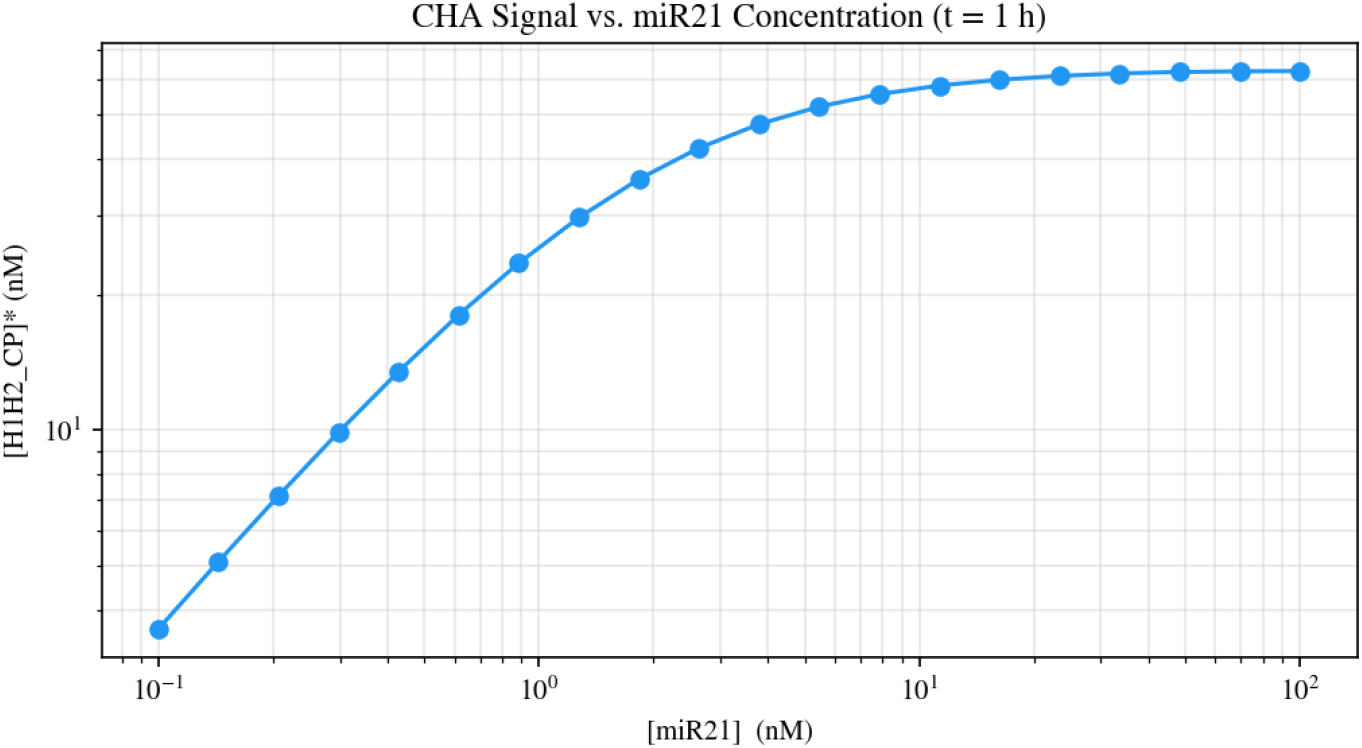
Steady-state reporter signal [*H*_1_: *H*_2_:CP]^∗^ as a function of the initial miR-21 concentration. The single-valued, monotone trace is consistent with the parameter-free uniqueness guarantee of the Deficiency One Theorem (*δ*_CHA_ = 1). Generated by `examples/04_cha_cascade.py`.

### 3.5 Goldbeter–Koshland switch: ultrasensitivity without bistability

The Goldbeter–Koshland (GK) covalent-modification cycle (Goldbeter and Koshland 1981; Ferrell and Ha 2014) is the canonical mechanism for *zero-order ultrasensitivity*. The full mass-action mechanism couples a kinase arm and a phosphatase arm,

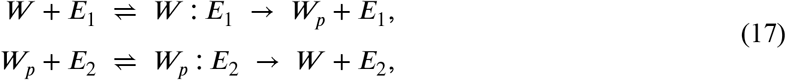

and possesses *m* = 6 species, *n* = 6 complexes, 𝓁 = 2 linkage classes, rank(**N**) = 3, so that *δ*_GK_ = 6 − 2 − 3 = 1. The network is *not* weakly reversible (the catalytic steps are irreversible), and each linkage class has per-LC deficiency 0 or 1 with at most one equal to 1: the Deficiency One Theorem therefore applies. The structural conclusion is that for *any* positive choice of *k*_on_, *k*_off_, 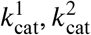 and *any* enzyme totals 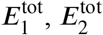, the mass-action system admits at most one positive equilibrium per stoichiometric compatibility class: **bistability is structurally impossible**.

The Goldbeter–Koshland quasi-steady-state (QSS) approximation (Goldbeter and Koshland 1981) predicts the fractional activation *y* ≔ [*W*_*p*_]^∗^/*W* ^tot^ as the unique root in (0, 1) of

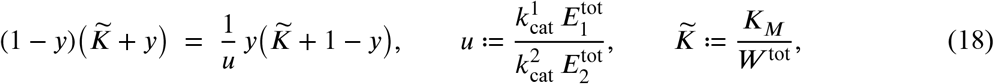

with *K*_*M*_ = (*k*_off_+*k*_cat_)/*k*_on_. In the deep zero-order regime *W* ^tot^/*K*_*M*_ ≫ 1, the curve *y*(*u*) becomes effectively step-like and the Hill coefficient *n*_*H*_ = log(81)/ log(EC_90_/EC_10_) diverges as *K*_*M*_ → 0. We use *W* ^tot^ = 1 M, 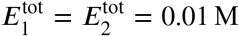, *k*_on_ = 10^4^ M^−1^ s^−1^, *k*_off_ = 0.1 s^−1^, *k*_cat_ = 1 s^−1^, giving *W* ^tot^/*K*_*M*_ ≈ 9000. Scanning 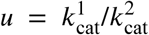 over more than two decades, `mantis-delta` finds exactly one steady state at every point (Fig. 5), confirming the D1T prediction. The numerical fractional activation agrees with (Equation 18) to better than 1% across the full scan, and the effective Hill coefficient at half-maximum exceeds 2 × 10^3^, illustrating the digital character of the switch.

**Figure 5:**
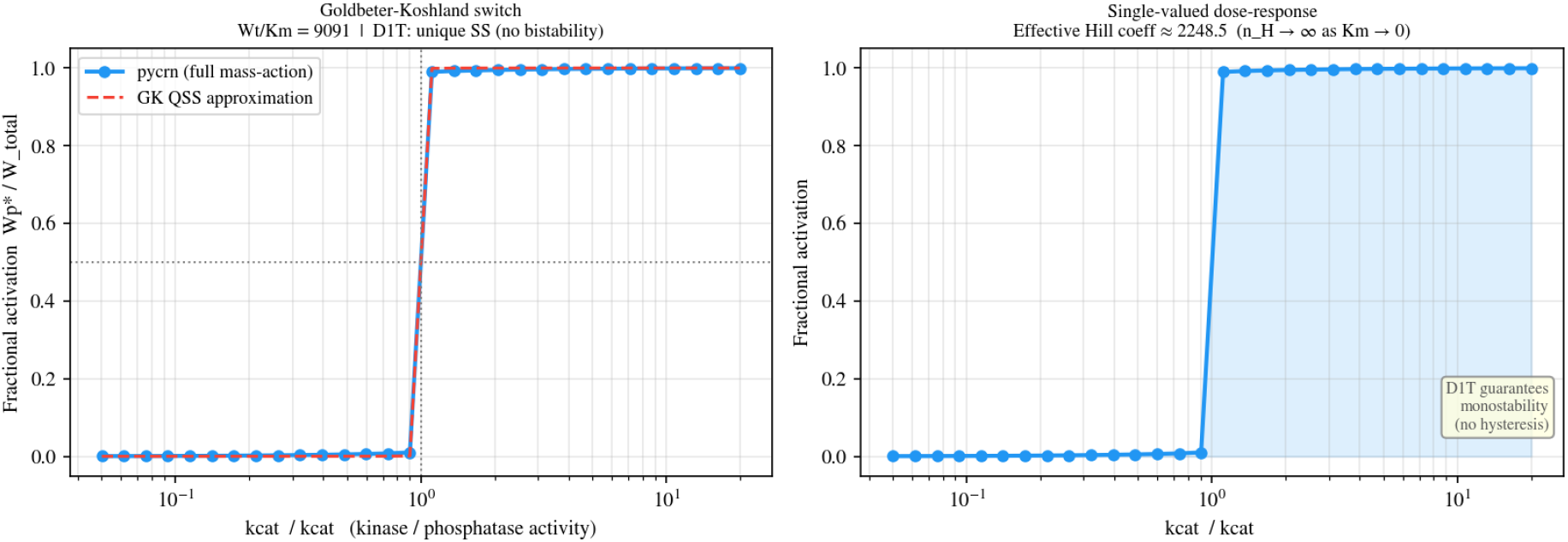
Goldbeter–Koshland zero-order ultrasensitivity switch. Left: fractional activation *y* = [*W*_*p*_]^∗^/*W* ^tot^ from `mantis-delta`’s full mass-action solution (markers) overlaid on the QSS prediction (Equation 18) (dashed). Right: the same data plotted alone, with the effective Hill coefficient annotated. The Deficiency One Theorem guarantees the single-valued nature of the curve. Generated by `examples/06_goldbeter_koshland.py`.

### 3.6 Reaction graph visualisation

For pedagogical exposition and debugging, `mantis-delta` exposes a NetworkX-based graph layout via `CRNetwork.draw()`. Figure 6 shows the complex reaction graph for the CHA network of §3.4; nodes are complexes, directed edges are reactions, and the four weakly connected components correspond to the four linkage classes 𝓁 = 4 that enter the deficiency formula (Equation 6).

**Figure 6:**
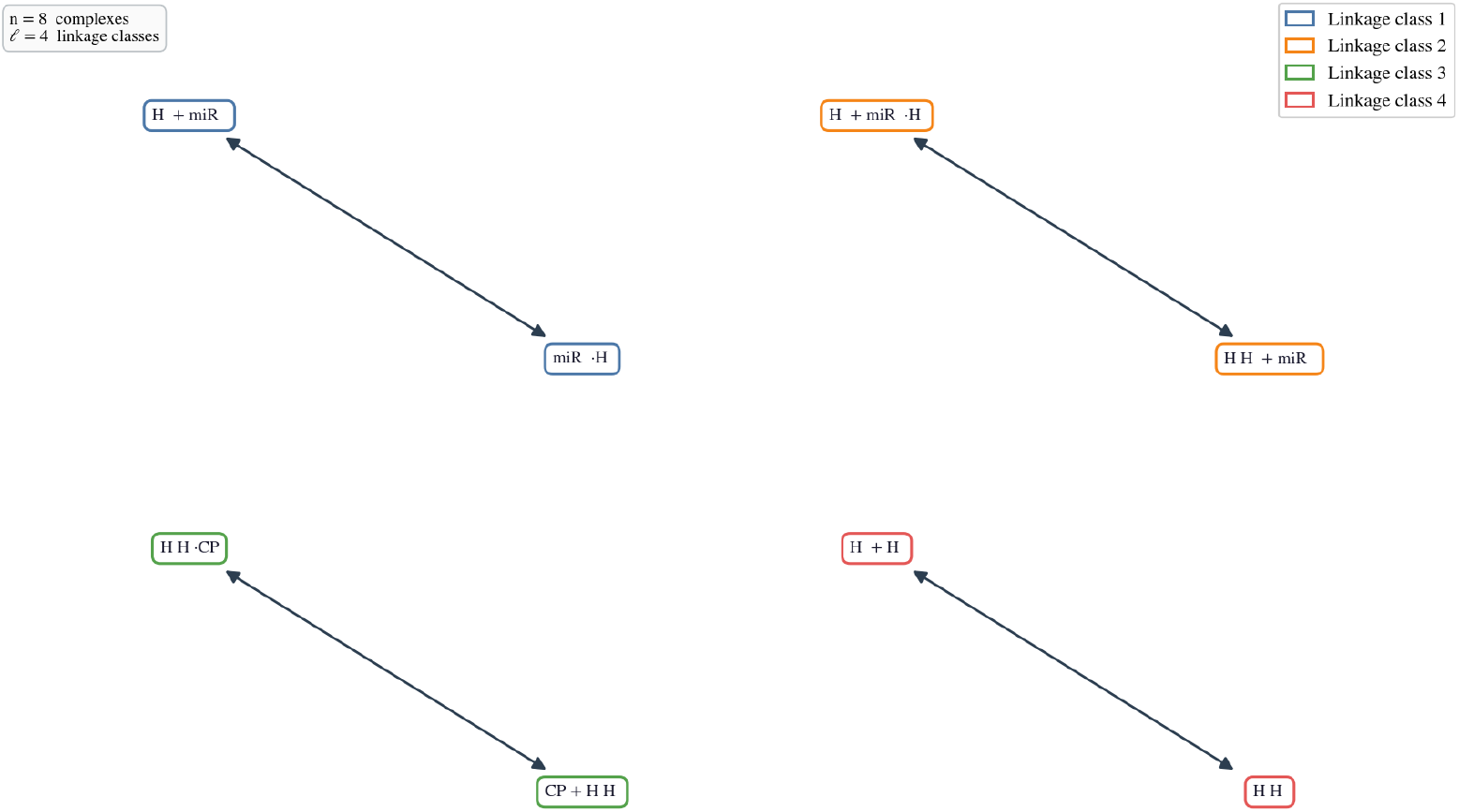
Complex reaction graph for the CHA miR-21 biosensor; nodes are complexes (e.g. miR21 + *H*_1_), directed edges are reactions. Four weakly connected components (𝓁 = 4) and eight complexes (*n* = 8) on a stoichiometric subspace of dimension *s* = 3 give *δ* = 8 − 4 − 3 = 1.

## 4 Discussion

We have presented `mantis-delta`, a lightweight Python implementation of Feinberg-style chemical reaction network theory tightly integrated with SymPy (Meurer et al. 2017) for symbolic ODE construction and SciPy (Virtanen et al. 2020) for numerical equilibration. The combination of parameter-free structural certification (DZT, D1T) with a hybrid integration/algebraic solver capable of locating both stable and unstable fixed points addresses a long-standing gap between the mathematical theory and applied practice: practitioners can now state, programmatically and prior to any wet-lab experiment, whether a given network is *structurally* prohibited from exhibiting bistability or oscillation, and then back-test that prediction across the whole physically plausible parameter regime by way of (Equation 3) and (Equation 8). The six benchmarks of §3 exercise the principal capabilities of the library: in all cases the CRNT-predicted qualitative behaviour is recovered with a solver residual below 10^−6^ M s^−1^, and the Goldbeter–Koshland dose-response curve agrees with the closed-form QSS approximation to within 1% over the entire 400× activity scan.

Compared with the foundational Chemical Reaction Network Toolbox (Ji et al. 2011) and the constraint-programming approach of Soliman (2012), `mantis-delta` offers (i) an idiomatic Python API tailored to the modern scientific stack (Harris et al. 2020; Virtanen et al. 2020), (ii) first-class support for chemostatted (open) networks in the spirit of Polettini and Esposito (2014), including the algebraic location of Hopf-unstable fixed points that forward integration cannot reach, and (iii) symbolic ODE/Jacobian expressions ready for downstream pipelines such as parameter inference, sensitivity analysis, or thermodynamic constraint checking (Ederer and Gilles 2007).

Several limitations remain. The current solver does not implement higher deficiency theorems (e.g. deficiency-two or higher-deficiency results) nor the recent toric-balanced equilibria framework of Craciun et al. (2009); the global attractor conjecture (Anderson 2011) is not invoked. Bifurcation scans are presently limited to one rate constant at a time, and continuation-style tracking of unstable branches across folds, as provided by XPPAUT (Ermentrout 2002) or AUTO, is not (yet) implemented. Model-reduction techniques (Rao et al. 2014) and robustness analysis (Shinar and Feinberg 2010) are natural next steps, as is integration with SBML import/export.

In conclusion, `mantis-delta` provides an accessible, well-tested, and mathematically rigorous entry point into chemical reaction network theory for the broader Python scientific community. By coupling Feinberg’s structural theorems to symbolic and numerical tools, it lowers the barrier between formal CRNT and practical model analysis, with applications spanning DNA computing (Yin et al. 2008; Zadeh et al. 2011), enzymatic signal transduction (Goldbeter and Koshland 1981; Ferrell and Ha 2014), and synthetic-circuit design (Ang et al. 2013).

## Data and code availability

All source code, examples, the 93-test pytest suite, and scripts to regenerate every figure in this paper are openly available under the MIT License at https://github.com/emiliovenegas/mantis-delta. The version used to prepare this manuscript is tagged `v0.1.0`.

## Author contributions

E.V. designed the library, implemented the code, performed all numerical experiments, and wrote the manuscript.

## Acknowledgements

The author acknowledges the open-source scientific Python ecosystem (Harris et al. 2020; Virtanen et al. 2020; Meurer et al. 2017; Hagberg et al. 2008) without which this work would not have been possible.

## Notes

### Competing Interest Statement

The authors have declared no competing interest.

